# Comparative Analysis of Non-linear Behavior with Power Spectral Intensity Response Between Normal and Epileptic EEG Signals

**DOI:** 10.1101/173518

**Authors:** Budhaditya Ghosh, Sourya Sengupta, Sayan Nag, Sayan Biswas, Shankha Sanyal

## Abstract

Epilepsy is a neurological condition which affects the nervous system. It is a general term used for a group of disorders in which nerve cells of the brain discharge anomalous electrical impulses from time to time, causing a temporary malfunction of the other nerve cells of the brain.EEG signal provides an important cue for diagnosis and interpretation related to prognosis of epilepsy. In this work we envisage to provide novel tool which can be used to detect the prognosis of epileptic disorder by comparing linear and non-linear modalities of EEG analysis – conventionally used Power spectral analysis and a robust non linear method – Detrended Fluctuation Analysis (DFA). Publicly available dataset is used for this work consisting of 100 normal patients’ EEG data as control group and 100 epileptic patients’ EEG data for comparison. Response for different frequency bands (alpha, theta, beta) of the EEG spectrum have been analyzed using Detrended Fluctuation Analysis (DFA) and Power Spectral Intensity (PSI). The comparison of the DFA scaling exponent with the spectral power data is calculated for all the 3 different frequency bands of EEG signal provide new and interesting results which have been discussed in detail.

## I. INTRODUCTION

EEG or Electroencephalography is an aperiodic time vs. amplitude plot in of electrical activity of the cerebral cortex nerve. In the signal which information regarding the activity of the cerebral cortex nerve is stored. [1]Many diseases can be diagnosed from the basic EEG responses of the patients. Epilepsy is one of the many diseases which can be diagnosed and analyzed with the help of EEG signal patterns and variations. [2][3]As the actual cause of epileptic seizure is not an unique one, so it can not be uniquely determined but nevertheless it can be said that sudden and random seizure discharge of various brain neurons that temporarily hampers functions of the brain may lead to it. The alarming fact is that globally there are 2.4 million new cases of epilepsy each year and this disease can be fatal in serious cases.

Epilepsy monitoring through different computer aided techniques is an important area of research nowadays. To monitor or recognize or distinguish epileptic EEG signals from the normal brain responses feature selection must be an important step in the total procedure. Features can be linear and non-linear in nature. As EEG signals are basically inherent so it is expected that the non-linear fractal analysis of EEG signals would lead to a good distinguishable approach or behavior of EEG signals. Main motivation behind this work is to study the behavior of two completely different parameters on the same EEG dataset and to find the correlation between them. No previous work has been done regarding this EEG analysis of normal and epileptic patient with two contrasting parametric approach.

Range of frequency of EEG signals can be divided as : (i) delta (δ) 0-4Hz, (ii) theta (θ) 4–8 Hz, (iii) alpha (α) 8- 13Hz and (iv) beta (β) 13-30 Hz.

In our analysis for the sake of convenience we have taken theta, alpha and beta range for our analysis purpose as this three are the major regions of the EEG signal. In this work we have compared non-linear and linear parametric behavior of EEG responses for epileptic patients and normal patients for above mentioned three frequency ranges.

A generic DFA algorithm was executed on the EEG signals which produced the hurst exponent for each EEG signals. The hurst exponent is essentially the non linear parameter. PSI (Power Spectral Intensity) served as a linear parameter in our work.

This type of analysis will pave the way for finer and detailed application of Brain-Computer Interaction.

A brief of the topics on which the different sections are concerned is given as follows. In section II information about EEG datasets used in our work is given. Section III and IV depicts the algorithms and corresponding mathematics of DFA and PSI .. Section V presents the detailed analysis of obtained result. Section V deals with conclusion and future research scope regarding this topic.

## II. EEG DATA

We have used a publicly available EEG time series database [4]. All the signals which have been used are taken from a pre amplifier system having 128-channel. The data was digitized with 12 bit resolution with a sampling rate of 173.61 samples per second. The database contains 5 sets of EEG signals which have been named as Set-A, Set-B, Set-C, Set-D and Set-E. Each of them has different significance [6]. Each of the dataset has EEG signals. All the signals have been taken from 100 single channel. Set A and set B of the total dataset have been taken from surface EEG recordings. Set A was for normal healthy patient with eyes open and set B was for normal healthy patients with eyes closed. For set C and D Signals were measured in seizure-free intervals from five patients. In the set C the signals were measured from the hippocampus formation of the opposite hemisphere of the brain. For set D it was in the epileptogenic zone. The last of the total dataset, set E has signals for seizure activity and these signals exhibit ictal activity.

We have chosen set A and set E for our analysis. We have tested the results upon set A and set E. Different type of EEG signals are shown in Fig 1.X axis is time(in Seconds) and Y axis is amplitude(in V)

**Fig 1:**
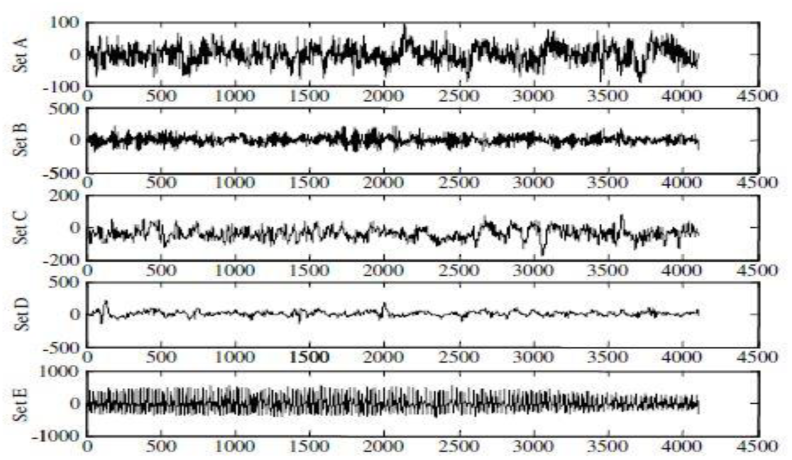
All 5 sets of EEG signals

## III. DETRENDED FLUCTUATION ANALYSIS

Detrended Fluctuation Analysis has proved to be benificial in understanding unique insights into neural organization. [5][8] DFA for a time series say {t1; t2; t3::::::tn} can be computed by following:

1. Another series T as [T(1); T(2); T(3)……T(N)], 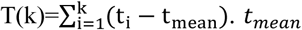 denotes mean of the points in the series t.
2. The series T is under interest. Series T is sliced into threads of length N. Each thread must contain this same number of element which is N. For each of the N element thread, a line is fit which signifies the trend in the thread. The fit is called Tn(k).
3. The detrending is [*T*(*k*) – *T_n_* (*k*)]^2^ which helps in calculation of RMS fluctuation. 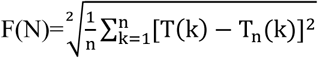 is called root mean square fluctuation.
4. F(n)∝ n^a^, a is expressed as the slope of logarithimic plot of log[F(n)] versus log(n).

Obtained a is the DFA value of a signal. It is called the DFA scaling exponent and it quantifies self-similarity and correlation properties of time series. As it suggests, a time series having has higher DFA scaling exponent is a symbolic quantification of presence of long range correlation. DFA quantifies complexity of using fractal property.

The scaling exponent a denotes the following:

1. a < 0.5 anti correlated
2. a = 0.5 white noise
3. a> 0.5 positive autocorrelation
4. a=1 1/f noise
5. a=1.5 Brownian noise

The DFA exponent can completely signify auto correlation of a signal. DFA technique was applied following the NBT algorithm used in Hardstone et.al [6]

**Fig 2:**
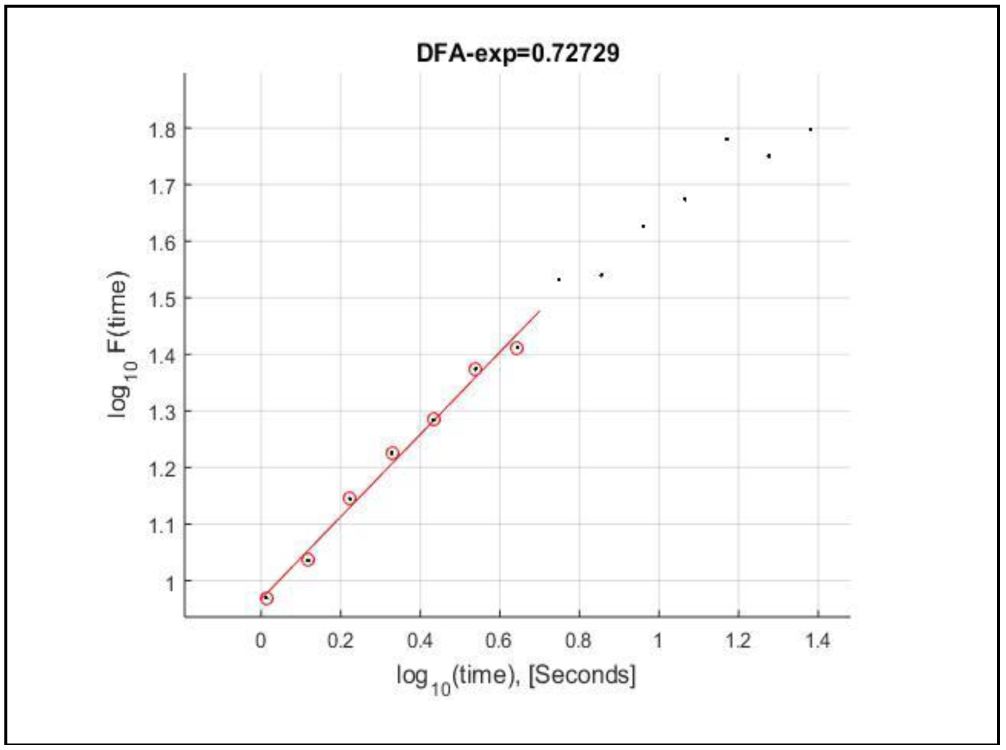
Sample plot for DFA of EEG signal

## IV. POWER SPECTRAL INTENSITY

The power spectrum of a time series data describes the distribution of power into frequency components composing that signal. [7]According to Fourier analysis any physical signal can be decomposed into a number of discrete frequencies, or a spectrum of frequencies over a continuous range.

To the time series data [x1,x,……,xN], we perform the Fast Fourier Transform (FFT) and the result obtained is denoted as [X1,X2,……,XN]. A continuous frequency band from flow to fup is sliced into K bins, which can be of equal width. We consider a vector band = [f1,f2,……,fK], such that the lower and the upper frequencies of the ithbin are fi and fi+1 respectively. The bins are used as θ (4-7 Hz), α (8-12 Hz) and β (12-30Hz). For these bins we have the band = [4, 7, 12, 30]. The Power Spectral Intensity (PSI) of the kth bin is evaluated as

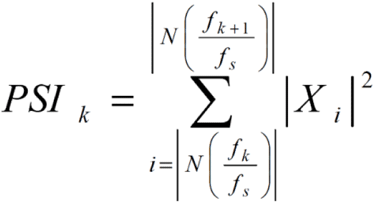

Where k=1, 2,….,K-1 where fs is the sampling rate, and N is the series length. We have converted the time series data into the frequency domain using Fast Fourier Transform (FFT) and the frequency descriptors of power bands, namely alpha beta and theta are extracted. This process is repeated for two sets (Set A and Set E) each of which contains 100 datasets. The values are normalized for easier comparison and plotted of graphs.

**Fig 3:**
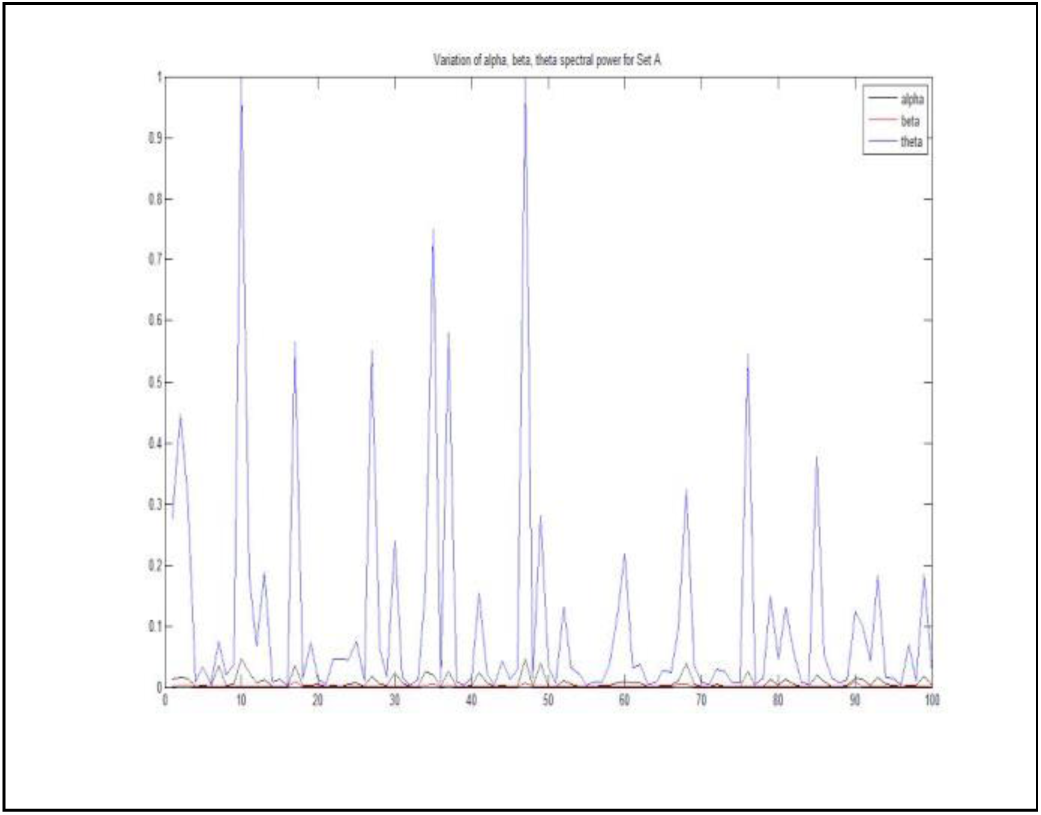
Sample plot for PSI of EEG signals

## V. RESULTS

Table 1 and Table 2 shows the DFA and PSI values of the total set A and E. Every set consists of 100 EEG signals. Hence the values shown here is the average value of all the 100 values obtained.

**TABLE 1:** DFA Values of both the sets

| SET | DFA values of two sets |  |  |
| --- | --- | --- | --- |
|  | *Alpha* | *Beta* | *Theta* |
| Set A | 0.667 | 0.539 | 0.690 |
| Set E | 0.631 | 0.605 | 0.660 |

**TABLE 2:** PSI Values of both the sets

| SET | PSI values of two sets |  |  |
| --- | --- | --- | --- |
|  | *Alpha* | *Beta* | *Theta* |
| Set A | 0.456 | 0.493 | 0.627 |
| Set E | 0.208 | 0.835 | 0.306 |

The graphs are plotted and discussed in the following figures below

Here in figure 4, figure 5 we have plotted bar graph and line graph of DFA values of Alpha, Beta and Theta Rhythms to get a overall comparison between a Normal man and an Epileptic patient. Here one can see the ranges of DFA values of each band, i.e. Alpha, Beta and theta are comparable. In Normal Condition, among Beta, Theta and Alpha, Theta takes the highest value and Beta takes the least and that kind of nature we can observe in Epileptic condition too. In Alpha Band, DFA for Normal condition is little bit higher than Epileptic condition and same as for Theta Band, Normal value is higher than Epileptic. But unlike them, In case of Beta rhythm, DFA value for Normal Man is less than the value for an Epileptic patient.

**Fig 4:**
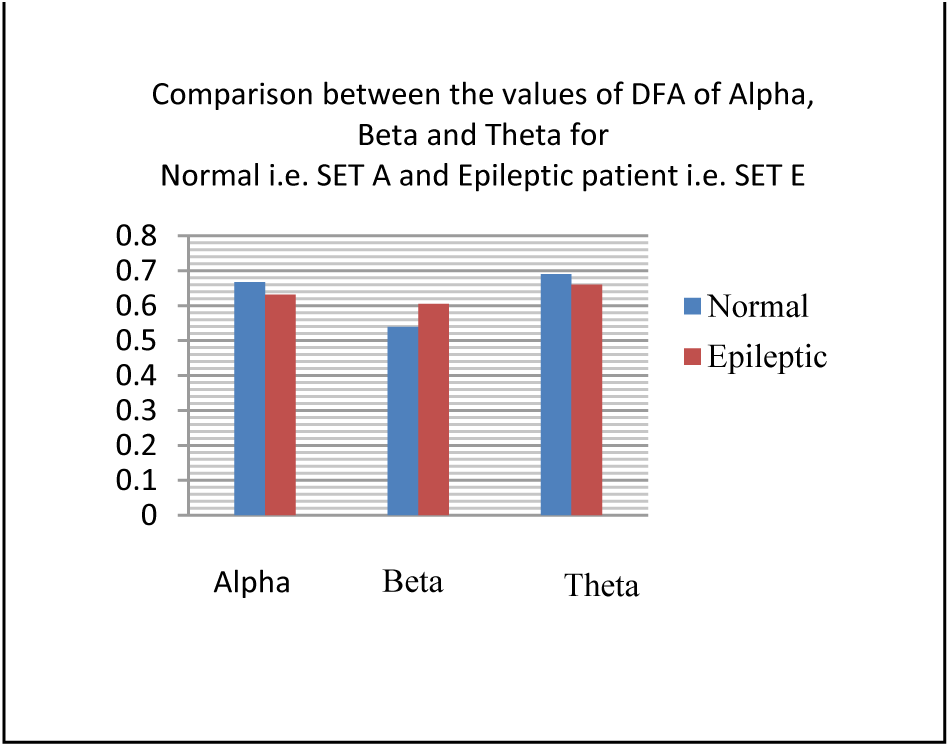
Bar Graph for Variation of DFA values of both set A and set E

**Fig 5:**
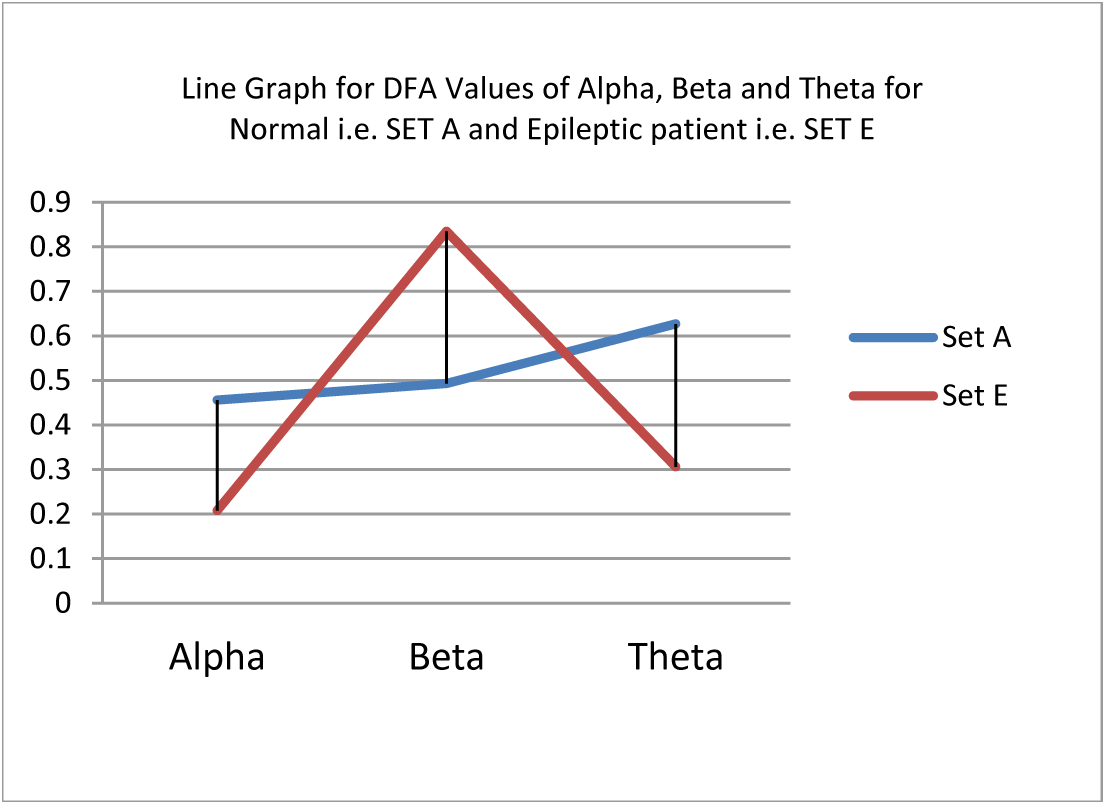
Line Graph for Variation of DFA values of both set A and set E

In the figure 6,figure 7 we have compared PSI values for Alpha band, Beta band and Theta band for both the datasets by plotting bar graph and linle graph. From this it can be easily observed that the variation of PSI value between normal and epileptic patients’ EEG signals are same as that of DFA values. The value of PSI is higher in Alpha and Theta region for normal patient than epilepsy patients. But for the Beta region the value for normal patient is lower than that of epileptic patients. The findings were pretty much same with DFA

**Fig 6:**
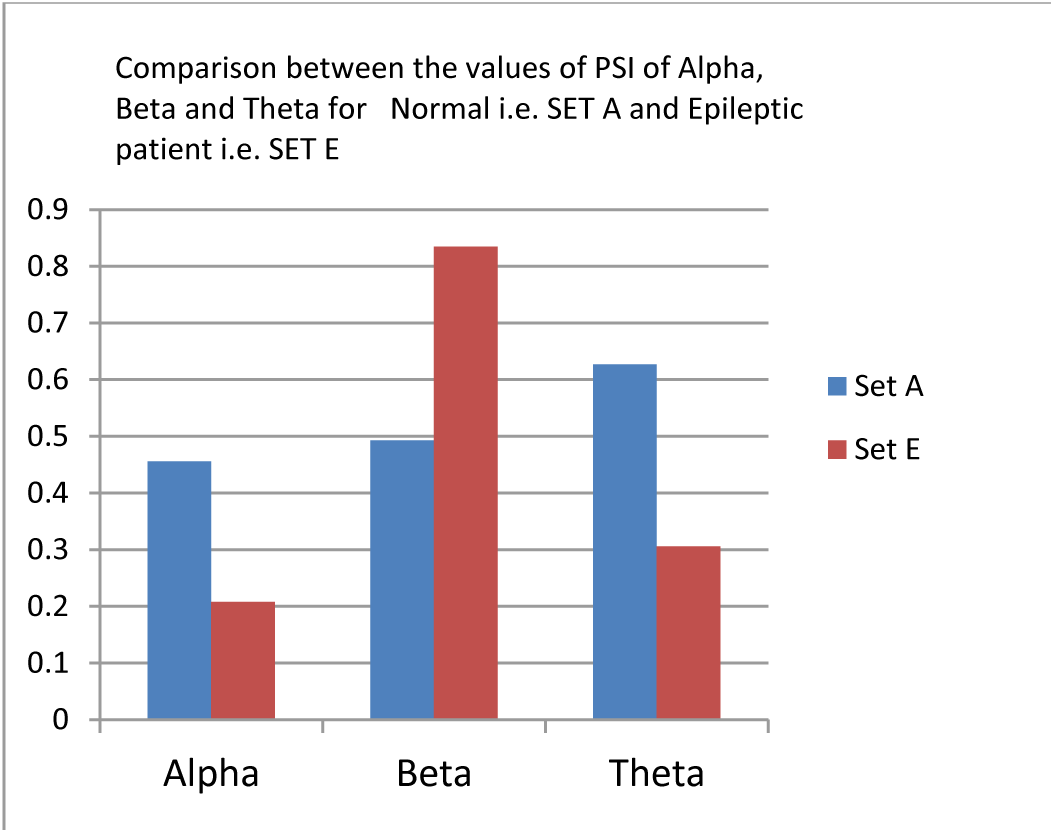
Bar Graph for Variation of PSI values of both set A and set E

**Fig 7:**
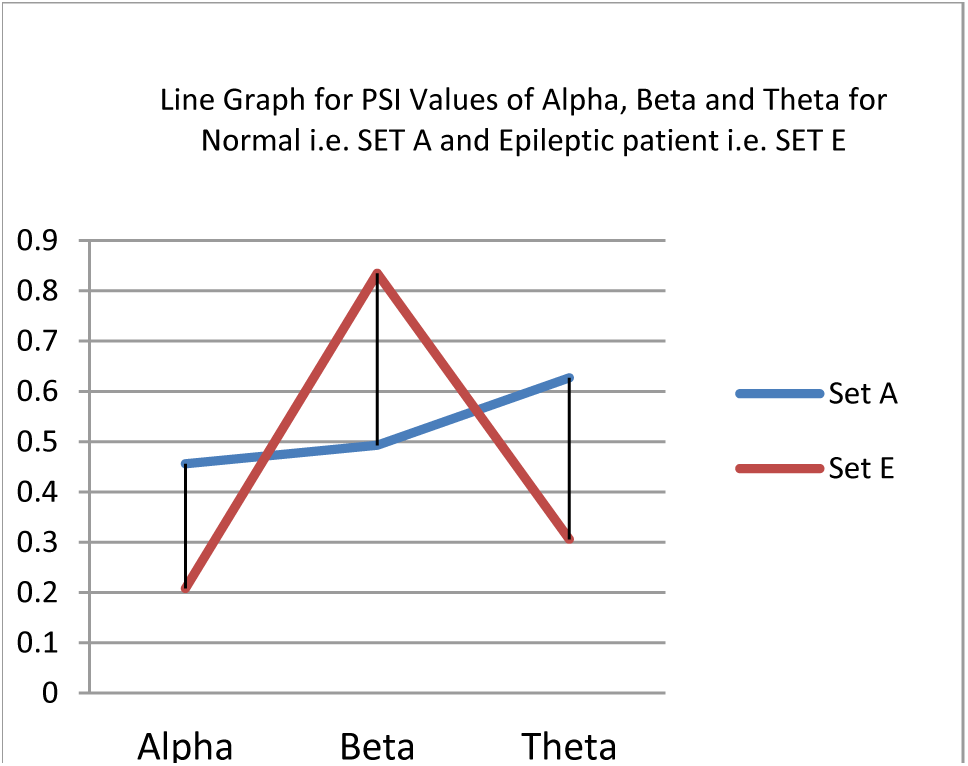
Line Graph for Variation of PSI values of both set A and set E

## VI. CONCLUSION AND FUTURE WORK

This work shows a detailed comparative analysis between DFA and PSI –two important parameters for statistical l\non linear and linear signal processing domain.

From this work, the major finding is quite interesting that the variation of linear and non-linear parameters for two completely different conditions i.e. normal and epilepsy is quite similar for different frequency region of EEG signal bands

The values of PSI and DFA for different frequency ranges are taken as average values from 100 normal patient signals and 100 epileptic signals.

Brain computer interface is gaining grounds in the research perspective as it allows endless possibilities in designing comprehensive interfacing. An parameter analysis of DFA and PSI as proposed in this work, could pave way for finer understanding of causalities occurring in the brain activity in normal patient and epileptic ones. Added to this BCI could be helped by this work as understanding brain activity is a initiation point in any BCI designing.

In this work we have just used two sets only for comparison however for future work purpose this results can be extended for all 5 sets.

